# Comparative transcriptomic analysis reveals conserved injury responses and divergent molecular responses underlying early regenerative competence following traumatic brain injury

**DOI:** 10.64898/2026.09.07.749916

**Authors:** Poulomi Ghosh, Subhra Prakash Hui

## Abstract

**Background:** Traumatic brain injury (TBI) is a leading cause of mortality and long-term neurological disability worldwide. Adult mammals have limited regenerative capacity following central nervous system (CNS) injury, but zebrafish have a remarkable regenerative potential, providing an opportunity to identify molecular mechanisms associated with successful CNS repair.

**Methods:** We have integrated publicly available acute-phase TBI transcriptomic datasets from human, mouse, rat, and zebrafish studies published up to April 2026. Differential expression data were combined using restricted maximum likelihood (REML)-based random-effects meta-analysis to account for biological and technical heterogeneity. Following ortholog mapping, we compared conserved injury responses with transcriptional programmes preferentially enhanced in zebrafish.

**Results:** The acute molecular response to TBI is largely conserved across vertebrate species with predominant activation of cell cycle regulation, chromosome segregation, and interferon-mediated immune signalling genes. A conserved injury-responsive gene set further highlighted enrichment of mitotic and JAK–STAT signalling. In contrast, 54 zebrafish-enhanced genes displayed preferential activation of wound healing, extracellular matrix remodelling, integrin-mediated signalling, and regulated immune responses. Protein–protein interaction analysis identified ***STAT1*, *STAT3*, *FN1*, *CD44*, *ANXA2*,** and ***AXL*** as central hub genes connecting these injury and repair-associated responses.

**Conclusions:** Our study suggest that successful CNS regeneration is not driven by unique injury pathways but rather by coordinated modulation of conserved injury responses and additional molecular programmes that promote tissue remodelling and repair. These regeneration-associated networks provide potential targets for future functional studies and may contribute to the development of regenerative therapeutic strategies for traumatic brain injury.

**Highlights:**

- Comparative transcriptomics reveals molecular programmes linked to CNS repair.
- Acute TBI elicits a conserved transcriptional response across vertebrates.
- Regenerative competence arises from coordinated immune and repair pathways.
- Zebrafish preferentially activate wound healing and ECM remodelling after TBI.
- *STAT1*, *STAT3*, *FN1* and *CD44* form central regenerative network hubs in zebrafish TBI.

**Graphical Abstract:** 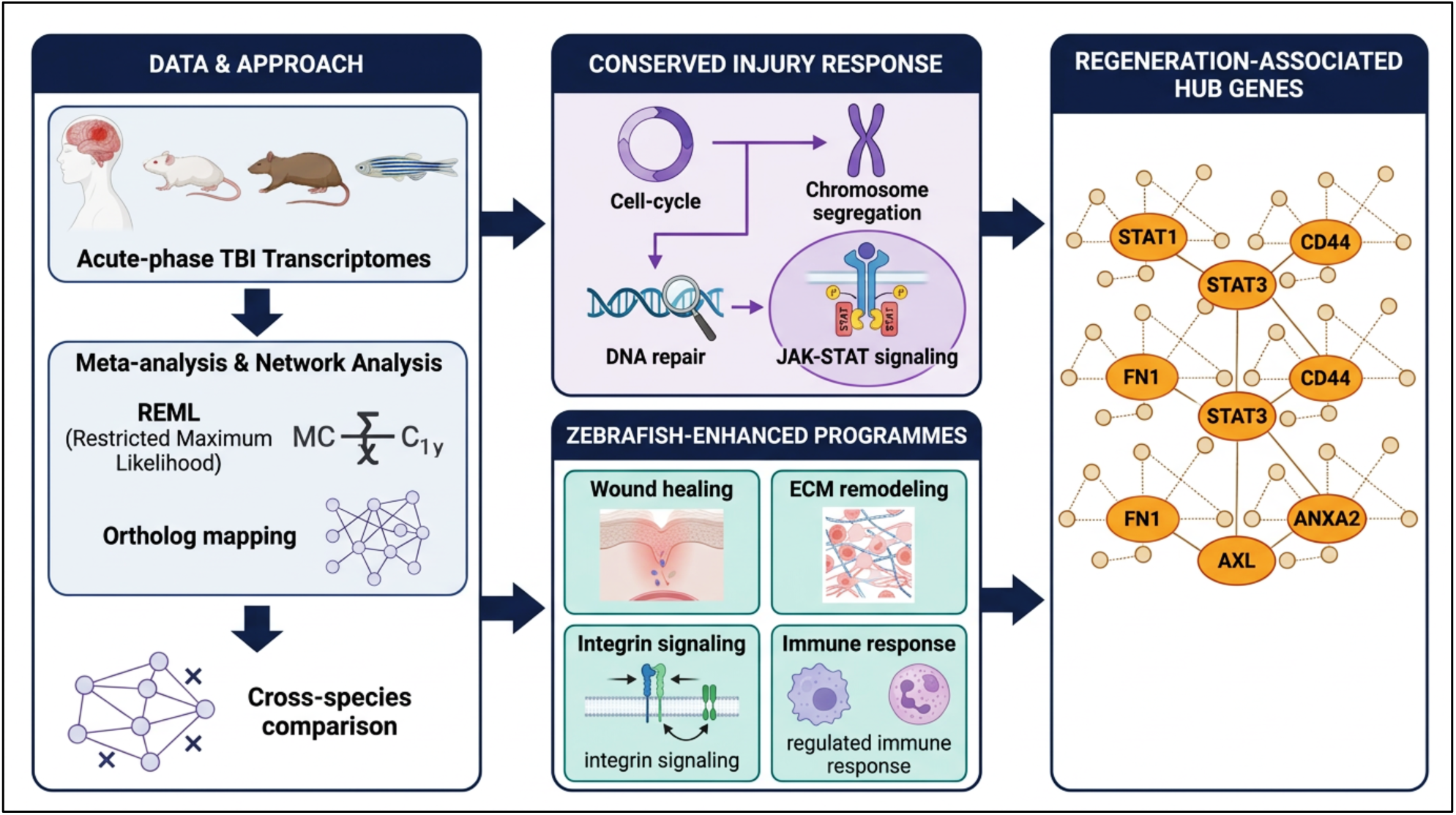

Cross-species meta-analysis of acute TBI transcriptomes (human, mouse, rat, zebrafish) reveals a conserved injury-response programme (cell-cycle, chromosome segregation, JAK– STAT signalling) alongside a zebrafish-specific regenerative programme (wound healing, ECM remodelling, integrin signalling). [Few parts of the graphical abstract created in BioRender. (https://BioRender.com)]

## 1. Introduction

Traumatic brain injury (TBI), sustained through falls, road-traffic collisions, and military combat, remains a leading cause of mortality and long-term neurological disability worldwide [1–6]. It affects people of every age group and imposes a substantial socioeconomic burden on health-care systems [7–9]. After the primary mechanical insult, a cascade of secondary injury processes ensues, encompassing blood–brain barrier disruption, neuroinflammation, excitotoxicity, oxidative stress, and progressive neuronal death [10,11], leaving many survivors with persistent cognitive, sensory, and motor impairments [4,12,13]. Despite advances in neurocritical care, therapies that meaningfully restore damaged neural circuits and functional recovery after TBI remain limited [14–16], largely because the adult mammalian central nervous system (CNS) has only a limited capacity for regeneration: neuronal replacement, axonal regrowth, and circuit reconstruction in adult mammals fall far short of what is required to compensate for injury-induced tissue loss [17–20]. Defining the endogenous repair mechanisms that operate after TBI is therefore essential for developing regenerative therapies.

Lower vertebrates such as the zebrafish (*Danio rerio*) stand in sharp contrast, possessing a profound capacity to regenerate the central nervous system (CNS) [21–25]. Following injury, the mammalian brain mounts prolonged neuroinflammation, reactive gliosis, restrained extracellular matrix remodelling, and limited neuronal replacement, a combination that foreclose successful regeneration [26–29]. The zebrafish brain, by contrast, is first sealed with scar tissue as inflammation subsides, but this scar is subsequently resolved and replaced by new, functional neurons generated from proliferating CNS-resident progenitor cells [30–34]. These divergent outcomes indicate that successful CNS regeneration is governed by distinct, evolutionarily conserved molecular pathways, and position comparative transcriptomic analysis across regeneration-competent and regeneration-limited vertebrates as a direct route to understand the genetic networks and biological pathways underlying successful CNS repair.

Transcriptomic technologies, including microarray profiling, bulk RNA sequencing (RNA-seq), and single-cell RNA sequencing (scRNA-seq), have transformed our capacity to dissect the molecular mechanisms of TBI [35–37], identifying genes and signalling pathways governing immune activation, inflammation, cell death, neurogenesis, angiogenesis, extracellular matrix remodelling, and tissue repair [38–41]. Yet most of these studies have examined single species, specific experimental models, isolated cell populations, or individual transcriptomic platforms, and substantial heterogeneity in injury models, sampling time points, sequencing technologies, and analytical pipelines has persistently obstructed the identification of robust molecular signatures reliably associated with regeneration, precluding direct comparison across studies.

Comparative transcriptomics across species with contrasting regenerative capacity offers a powerful strategy for resolving conserved injury responses from regeneration-associated molecular programmes, yet no study, to our knowledge, has undertaken an integrative analysis spanning the full range of publicly available TBI transcriptomic datasets across regeneration-limited and regeneration-competent vertebrates. Closing this gap requires an analytical framework capable of harmonising heterogeneous datasets generated across transcriptome platforms, acute post-injury time points, and cell-type-specific expression profiles while preserving biologically meaningful signal, thereby exposing molecular pathways that are consistently associated with successful CNS regeneration across independent studies.

Here, we systematically curated every publicly available acute-phase TBI transcriptomic dataset in human, mouse, rat, and zebrafish reported through 16 April 2026. To resolve technical and biological heterogeneity across these datasets, we applied a random-effects meta-analysis using the restricted maximum likelihood (REML) estimator, integrating repeated gene-level expression estimates arising from different acute-phase time points and cell-type-specific transcriptomic profiles into harmonised, species-specific expression datasets. Following ortholog mapping, we identified conserved, divergent, and regeneration-associated molecular programmes through extensive cross-species transcriptome comparison, functional enrichment analysis, and network-based biological interpretation. By integrating heterogeneous transcriptomic evidence across vertebrates with contrasting regenerative capacities, our study establishes a comprehensive framework for understanding the molecular underpinnings of CNS regeneration after TBI and nominates candidate pathways for future regenerative therapeutic strategies.

## 2. Methods

### 2.1 Literature curation and study selection

We performed a systematic literature search to identify the publicly accessible traumatic brain injury transcriptomic studies in human, mice, rat and zebrafish. We conducted the search using the NCBI PubMed and the NCBI Gene Expression Omnibus (GEO) database for studies published till 16^th^ April 2026. The search terms included “Traumatic Brain Injury”, “Transcriptomics”, “TBI”, “Transcriptome” and the respective species names.

We included only those studies which were generated using genome-wide transcriptomic data from species of wild-type strain, acute TBI brain samples (till 72 hours post injury(hpi)), included an uninjured group (control), no drugs were administered before or after TBI and provided abundant information for differential expression analysis. Studies were omitted out which reported use of transgenic line for sequencing, only targeted gene expression, lacked suitable control groups, certain chemicals or drugs were used or comprised of insufficient data for downstream analyses.

### 2.2 Data processing and integration of transcriptomic data

The eligible datasets were then grouped according to species and transcriptomic platform. Official gene symbols were used as gene identifiers and differential expression measurements (log₂ fold change and associated statistical significance) were curated or calculated for each dataset. The differential gene expression measurements were calculated using Seurat R package (version 5.4.0) and DESeq2 R package (version 1.46.0). As our included studies differ in sequencing platform, experimental design, and sampling time, each dataset was analysed independently before integration. To reduce the effects of study heterogeneity, repeated expression measurements of the same gene were amalgamated using a random-effects meta-analysis with the restricted maximum likelihood (REML) estimator. Differential gene expression estimates from individual cell populations were integrated to obtain a single gene-level estimate for each species in case of single-cell RNA-seq datasets. And for independent bulk RNA-seq or microarray studies and multiple acute post-injury time points, a similar estimator was applied to combine repeated measurements of the same gene to produce one representative expression value per gene for each species.

### 2.3 Mapping of orthologs and construction of comparative datasets

The orthologous genes shared among the four species were identified through the Ensembl BioMart database using the biomaRt R package (version 2.62.1). Two reference datasets were independently constructed, one with human gene symbols as the common reference and the other with mouse gene symbols as the common reference. The first dataset integrated gene expression profiles from all the four species. To maintain the robustness of the findings and reduce the potential bias from the limited availability of human datasets, the second one included mouse, rat and zebrafish transcriptomic profiles. Each meta-dataset comprised of log₂ fold change values and corresponding statistical significance for every orthologous gene and was used for downstream comparative transcriptomic analyses, functional enrichment, and identification of molecular pathways associated with regenerative competence following TBI.

### 2.4 Differential Gene Expression Analysis

We performed differential gene expression analysis independently for each transcriptomic dataset following species-specific pre-processing and REML-based integration, as described above. Differentially expressed genes (DEGs) were screened and retained for subsequent analysis using the criteria: log₂ fold change ≥ 1 and adjusted P value < 0.05.

Orthologous genes from the human-reference dataset, which included transcriptomic profiles from human, mouse, rat, and zebrafish were incorporated for the primary analysis. Consecutively, to estimate the robustness of the findings, a similar analytical workflow was applied to the independently constructed mouse-reference dataset comprising mouse, rat, and zebrafish. The numbers of significantly upregulated and downregulated genes were established for each species, and the overlap of orthologous DEGs across species was visualized using UpSet analysis.

### 2.5 Comparative transcriptomic analysis

The integrated orthologous gene expression profiles were compared between human and zebrafish, mouse and zebrafish, and rat and zebrafish using DEGs with an absolute log₂ fold change of at least 1. The genes were categorized as conserved upregulated, conserved downregulated, or oppositely regulated according to their direction of regulation between the regeneration-limited and regeneration competent vertebrates. Pairwise scatter plots were created to visualize the regulatory relationships among the species.

Thereafter, to evaluate the global expression patterns among conserved genes, hierarchical clustering was performed using scaled log₂ fold change values from all four species. Heatmaps were generated using the ComplexHeatmap package (version 2.22.0) in R, with row-wise Z-score normalization and unsupervised clustering of both genes and species to identify similarities in transcriptional responses following acute TBI among the vertebrates.

### 2.6 Identification of conserved injury-responsive genes

The conserved transcriptional responses following acute TBI were identified by filtering DEGs from the integrated comparative transcriptomic dataset based on differential expression in zebrafish and mammalian species. Genes exhibiting significant upregulation in zebrafish (log₂ fold change ≥ 1and adjusted *P* < 0.05) together with significant upregulation in at least one mammalian species (human, mouse, or rat) using the same significance thresholds were selected for downstream analyses. Human gene symbols were used as the reference identifiers for the primary analysis following ortholog mapping.

#### 2.6.1 Gene Ontology enrichment analysis of conserved injury-responsive genes

Functional enrichment analysis was performed using the clusterProfiler package (version 4.14.6) in R. Gene Ontology (GO) Biological Process enrichment was carried out using the filtered conserved gene set against the human genome annotation database using the org.Hs.eg.db (version 3.20.0) in R. Over-represented biological processes were identified using the Benjamini–Hochberg method for multiple-testing correction, and significantly enriched GO terms were retained using an adjusted *P*-value threshold of 0.05 Representative GO terms were visualized as bubble plots, where dot size represents the number of genes contributing to each biological process, colour indicates the adjusted *P*-value, and the x-axis denotes the gene ratio.

#### 2.6.2 Hierarchical clustering of conserved genes

The expression patterns of the conserved upregulated DEGs were visualized using heatmaps generated with the **ComplexHeatmap** package (version 2.22.0) in R. Log₂ fold-change values from human, mouse, rat, and zebrafish were standardized across each gene using row-wise Z-score transformation to facilitate comparison of relative expression patterns among species. Hierarchical clustering was performed using Euclidean distance and complete linkage for both genes and species.

### 2.7 Identification of zebrafish-enhanced genes

Candidate regeneration-associated genes were identified by using the orthologous gene expression profiles from the human-reference master dataset. Genes were defined as **zebrafish-enhanced** if they satisfied the following criteria: (i) significant upregulation in zebrafish (log₂FC ≥ 1 and adjusted *P* < 0.05), and (ii) a higher log₂ fold change in zebrafish than in human, mouse, and rat. This approach enabled the identification of genes displaying a preferential transcriptional response in the regeneration-competent species following acute TBI.

#### 2.7.1 Gene Ontology enrichment analysis of zebrafish-enhanced genes

Functional enrichment analysis of the zebrafish-enhanced genes was performed using the clusterProfiler package (version 4.14.6) in R. Gene symbols were converted to Entrez Gene IDs using the org.Hs.eg.db annotation database (version 3.20.0). Gene Ontology (GO) Biological Process enrichment analysis was subsequently performed using the enrichGO function with the Benjamini– Hochberg method for multiple testing correction. Biological processes with an adjusted *P* value < 0.05 were considered significantly enriched. Representative GO terms were selected for visualization in a bubble plot based on their biological relevance, where dot size represents the number of genes contributing to each biological process, colour indicates the adjusted *P*-value, and the x-axis denotes the gene ratio.

#### 2.7.2 Protein–protein interaction network analysis

Protein–protein interaction analysis was performed using the **STRING** database (Homo sapiens; version 12.0) to investigate functional associations among the zebrafish-enhanced genes.

#### 2.7.3 Hub gene identification

Network topology analysis was carried out by determining the degree centrality for each node in the PPI network via the **igraph** package (version 2.2.2) in R. Ranking of genes was done based on the degree value of the genes, while top ten genes with highest degree centrality were identified as hub genes. Degree centrality was chosen as one of the measures of network importance since it quantifies the number of direct interactions related to each protein in the interaction network.

## 3. Results

### 3.1 Systematic curation and integration of acute TBI transcriptomic datasets

To identify the molecular responses associated with CNS regeneration after TBI, we systematically curated all publicly available TBI transcriptomic datasets from human, mouse, rat, and zebrafish studies published through 16^th^ April 2026 in the NCBI PubMed and Gene Expression Omnibus (GEO) databases. This search yielded 749 literature reports, of which 35 datasets met our predefined eligibility criteria **(Fig. 1 and Supplementary Table S4)**. The final compendium, built from bulk RNA sequencing (RNA-seq), single-cell RNA sequencing (scRNA-seq), and microarray platforms, comprised 1 human, 17 mouse, 11 rat, and 6 zebrafish datasets.

**Fig. 1.**
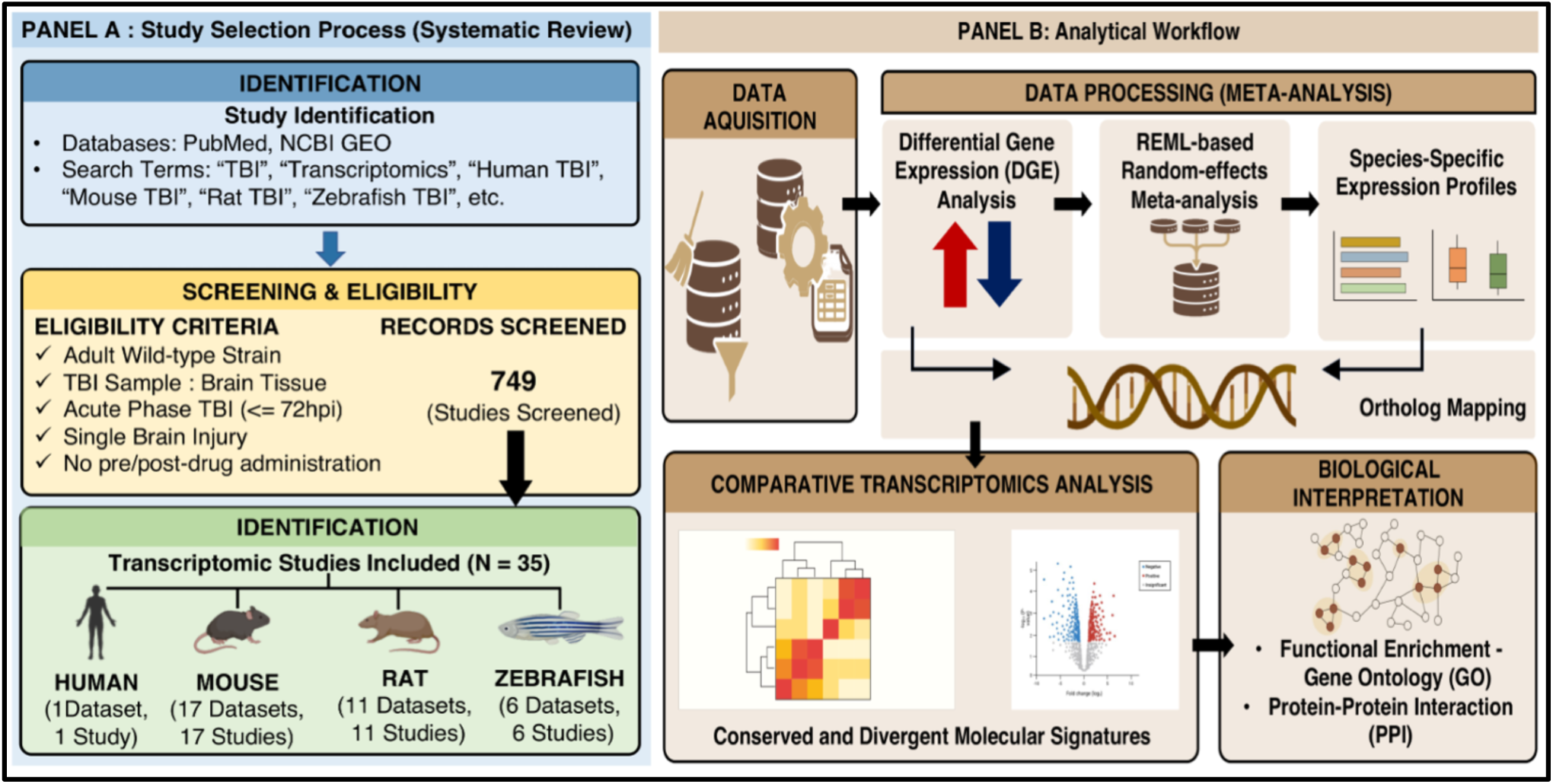
Legend: Methodological overview and analytical pipeline for multi-species comparative transcriptomic analysis. [Few parts of the illustration created in BioRender. (https://BioRender.com)]

Because the included studies differed in experimental design, sequencing platform, sampling time point, and biological source, we processed each dataset independently before integration, restricting our analysis to transcriptomic data generated from brain tissue during the acute phase of TBI to isolate the early injury response. Differential gene expression analysis was performed for each dataset, and restricted maximum likelihood (REML)-based random-effects meta-analysis combined repeated measurements of the same gene arising from different cell types, acute time points, or independent studies within a species **(Fig. 1)**, yielding a single representative expression estimate for each gene that accounted for variability between datasets.

To directly compare the regeneration-limited (human, mouse, rat) and regeneration-competent (zebrafish) species, we mapped orthologous genes and integrated two reference datasets: a human-reference dataset comprising human, mouse, rat, and zebrafish expression profiles **(Supplementary Table S5)**, and a mouse-reference dataset comprising mouse, rat, and zebrafish profiles **(Supplementary Table S6)**, constructed to safeguard against bias from the limited availability of human data. These two meta-datasets formed the basis of every subsequent comparative analysis of the molecular pathways underlying regenerative and regeneration-limited responses to acute TBI.

### 3.2 Global transcriptional responses to acute traumatic brain injury differ across vertebrate species

Differential gene expression profiles from the integrated human-reference dataset revealed significant transcriptional alterations in all four species following acute TBI, although the magnitude of response varied considerably: 27, 29, 59, and 69 differentially expressed genes (DEGs) were identified in human, mouse, rat, and zebrafish, respectively **(Fig. 2A)**.

**Fig. 2.**
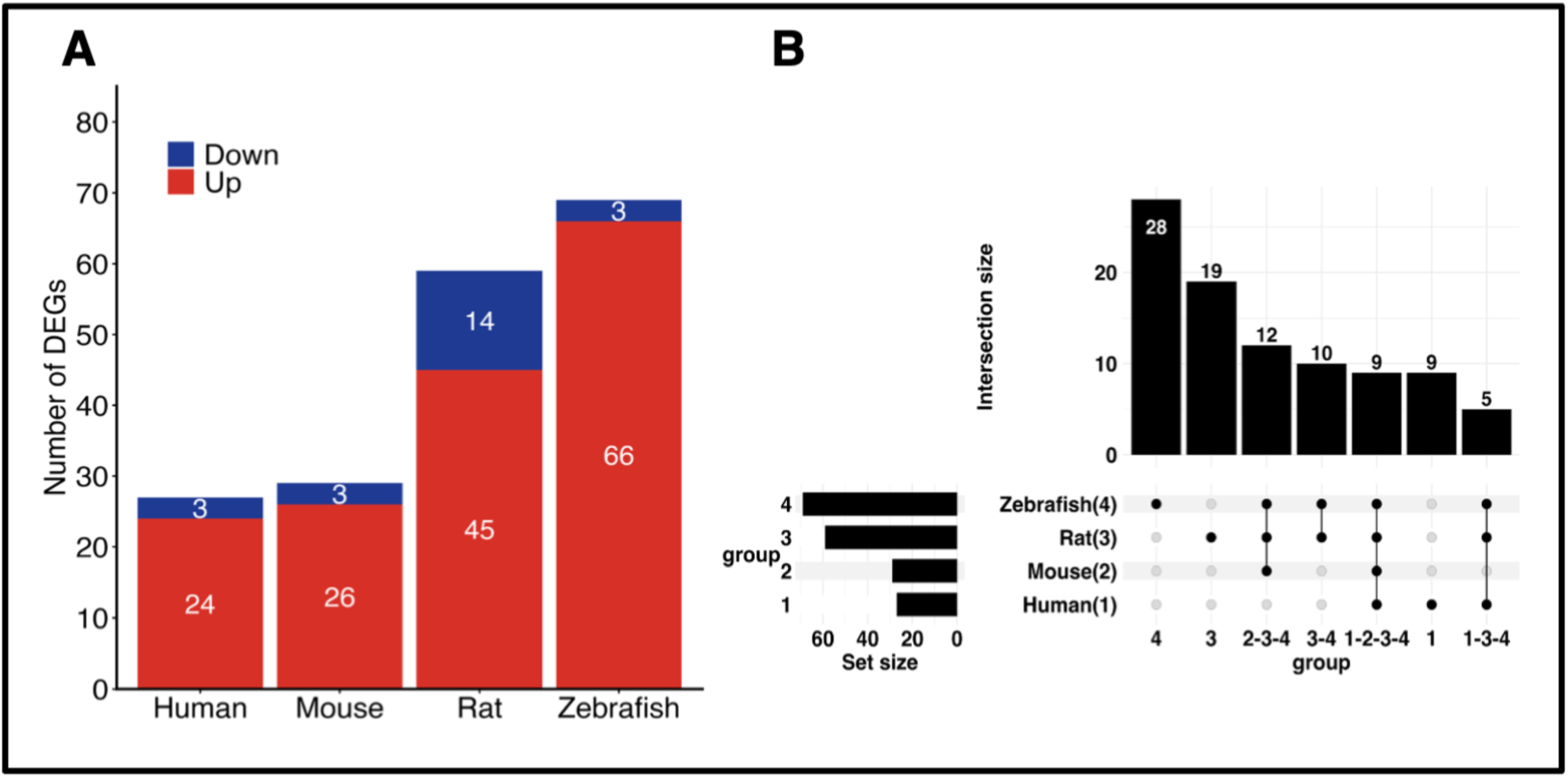
Legend: Global differential gene expression responses to acute traumatic brain injury across vertebrate species. (A) Number of significantly differentially expressed genes (DEGs) identified following acute traumatic brain injury in human, mouse, rat, and zebrafish after species-specific differential expression analysis and REML-based integration. Differentially expressed genes were defined using an absolute log₂ fold change (|log₂FC|) ≥ 1 and an adjusted *P* value < 0.05. Red bars represent upregulated genes, whereas blue bars represent downregulated genes. Zebrafish exhibited the largest transcriptional response, followed by rat, while human and mouse displayed comparatively fewer DEGs;(B) UpSet plot illustrating the overlap of orthologous DEGs among the four species using the human-reference comparative dataset. Horizontal bars indicate the total number of DEGs identified in each species (set size), whereas the vertical bars represent the number of genes shared among different combinations of species (intersection size). Filled black circles connected by vertical lines indicate the species included in each intersection. The majority of DEGs were species-specific or shared among a limited number of species, whereas only a relatively small subset of genes was conserved across multiple vertebrates, highlighting both conserved and species-specific transcriptional responses following acute TBI

Transcriptional activation dominated the acute TBI response across all vertebrates, with more upregulated than downregulated DEGs in every species. Zebrafish exhibited the largest transcriptional response, with 66 upregulated and 3 downregulated genes, followed by rat (45 upregulated, 14 downregulated); human and mouse showed more modest responses, with 24 and 26 upregulated genes, respectively, and only 3 downregulated genes each **(Fig. 2A)**. The magnitude and direction of transcriptional regulation after acute TBI therefore differ considerably across vertebrate species.

To assess the conservation of these responses, we compared orthologous DEGs across species using UpSet analysis **(Fig. 2B)**. Most differentially expressed genes were restricted to individual species or shared among only a limited subset, while a comparatively small set of DEGs was conserved across all four vertebrates. Notably, zebrafish displayed the largest proportion of species-specific DEGs, marking a transcriptional response distinct from that of the regeneration-limited species (human, mouse, rat). Although acute TBI elicits a common injury response, its underlying transcriptional programmes are predominantly species-dependent, motivating for subsequent comparative analyses to identify molecular signatures of regenerative competence.

This pattern was observed when the same analyses were repeated on the mouse-reference dataset: the overall distribution of differential gene expression and gene overlap remained similar **(Fig. S1)**, confirming that our principal findings do not depend on inclusion of the human dataset alone.

### 3.3 Comparative transcriptomic analysis divulges conserved and divergent molecular responses following acute traumatic brain injury

We next assessed the conservation of transcriptional responses between regeneration-limited vertebrates and regeneration-competent zebrafish by comparing orthologous genes with an absolute log₂ fold change ≥ 1. Pairwise analyses of the integrated four-species dataset revealed both conserved and species-specific patterns of gene regulation during the acute phase of TBI **(Fig. 3A–D)**.

**Fig. 3.**
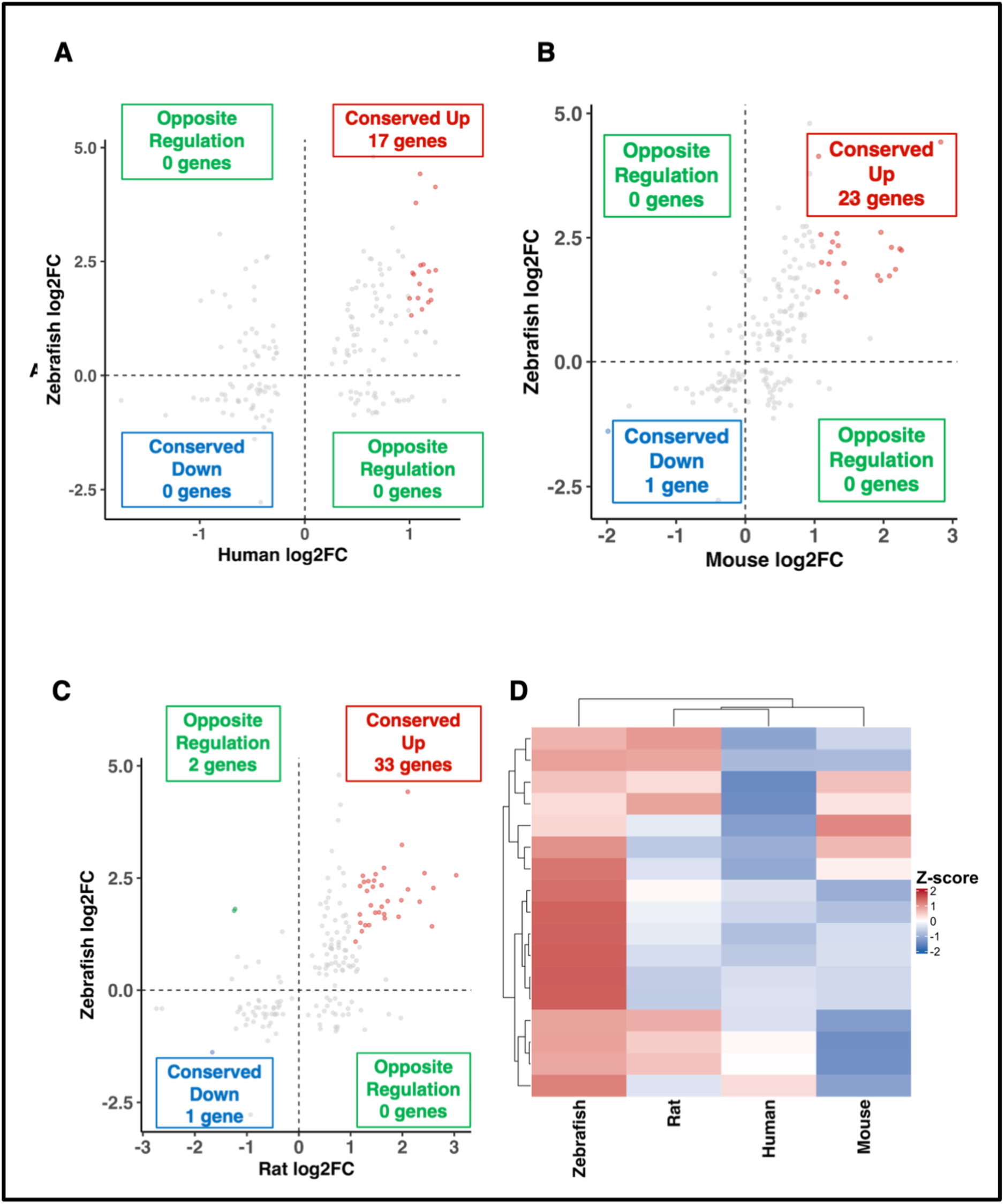
Legend: Comparative transcriptomic analysis identifies conserved and divergent transcriptional responses following acute traumatic brain injury across vertebrate species. (A–C) Pairwise comparison of orthologous gene expression between regeneration-limited vertebrates and the regeneration-competent zebrafish using the human-reference comparative dataset. Scatter plots compare log₂ fold-change values between (A) human and zebrafish, (B) mouse and zebrafish, and (C) rat and zebrafish. Each point represents a single orthologous gene. Dashed lines indicate the thresholds for differential expression (log₂FC = ±1 and 0). Genes significantly upregulated in both species (conserved upregulated) are shown in red, conserved downregulated genes in blue, oppositely regulated genes in green, and genes not meeting the differential expression criteria are shown in grey. The number of genes within each regulatory category is indicated in the corresponding quadrant;(D) Hierarchical clustering heatmap of conserved orthologous differentially expressed genes identified across the human-reference dataset. Log₂ fold-change values were standardized by row-wise Z-score transformation prior to clustering. Hierarchical clustering was performed using Euclidean distance and complete linkage for both genes and species. The heatmap demonstrates shared transcriptional responses among vertebrates while highlighting species-specific differences in the magnitude of gene expression following acute traumatic brain injury

Conserved upregulated genes constituted the dominant shared transcriptional response across all vertebrates during acute TBI: human and zebrafish shared 17 conserved upregulated DEGs, mouse and zebrafish shared 23, and rat and zebrafish shared 33. Conserved downregulated DEGs were essentially absent, none between human and zebrafish, and only one each in the mouse–zebrafish and rat–zebrafish comparisons and oppositely regulated genes were similarly rare, limited to two DEGs between rat and zebrafish **(Fig. 3A–C)**. The early response to TBI is therefore characterized predominantly by a conserved programme of transcriptional activation, with discordant regulation the exception rather than the rule.

Hierarchical clustering of the conserved orthologous DEGs further resolved the concordance and discordance in transcriptional responses across species during acute TBI **(Fig. 3D)**. A common set of injury-responsive genes was evident across vertebrates, but the magnitude of regulation differed by species, yielding distinct clustering patterns. Together, these results reveal both shared and species-specific molecular responses during the acute TBI phase, establishing the architecture from which we identify pathways associated with regenerative competence.

Consequently, we repeated the comparative analyses on the independently constructed mouse-reference dataset (mouse, rat, zebrafish) to test the robustness of these results **(Fig. S2)**. Conserved upregulated genes again dominated the transcriptional signature, with 182 conserved upregulated genes between mouse and zebrafish and 238 between rat and zebrafish, while conserved downregulated genes (three in each comparison) and oppositely regulated genes (11 in total) remained rare. Hierarchical clustering of the mouse-reference dataset reproduced the expression patterns of the primary four-species analysis, confirming the robustness of the comparative framework irrespective of the ortholog reference used.

### 3.4 Conserved injury-responsive genes reveal shared biological processes associated with acute traumatic brain injury

To pinpoint the biological processes that are consistently activated after acute TBI, we focused on DEGs significantly upregulated in zebrafish and in at least one mammalian species. This filtering strategy defined a conserved set of injury-responsive genes shared across regeneration-competent and regeneration-limited vertebrates, which were then subjected to Gene Ontology (GO) Biological Process enrichment analysis.

GO enrichment of the integrated human-reference dataset showed that conserved upregulated genes were chiefly associated with cell cycle progression and chromosome dynamics, chromosome segregation, sister chromatid segregation, nuclear division, and chromosome condensation **(Fig. 4A)** alongside enrichment of type II interferon-mediated signalling, cellular response to type II interferon, and JAK–STAT signalling-related pathways, indicating that conserved immune signalling accompanies the proliferative response during acute TBI. Cell-cycle regulation and immune activation therefore constitute the evolutionarily conserved components of early injury response.

**Fig. 4.**
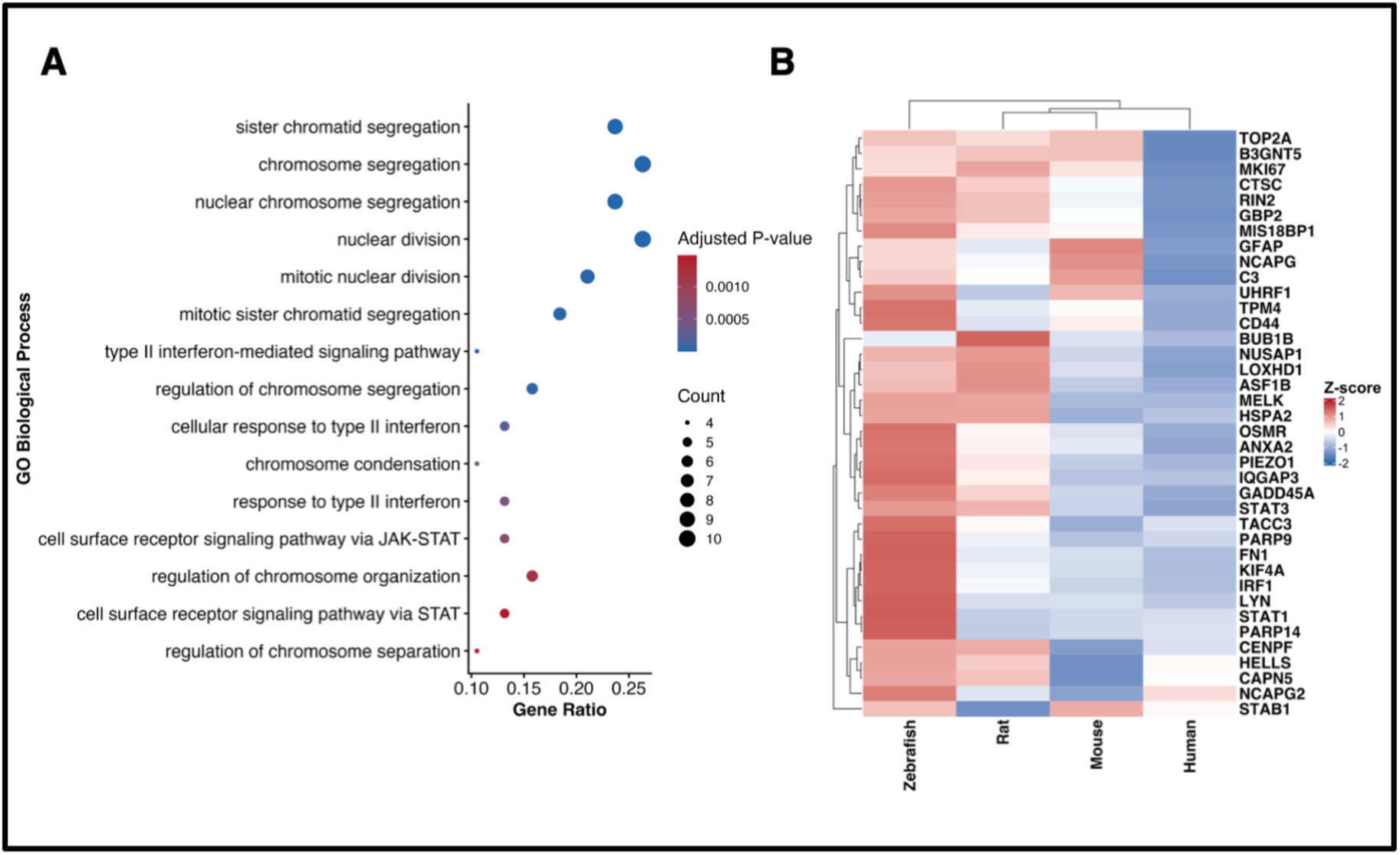
Legend: Conserved biological processes associated with the acute response to traumatic brain injury. (A) Gene Ontology (GO) Biological Process enrichment analysis of conserved upregulated genes identified using the human-reference ortholog dataset. Dot size represents the number of genes contributing to each enriched biological process, while colour denotes the adjusted *P*-value. The x-axis indicates the gene ratio for each GO term; (B) Hierarchical clustering heatmap showing the relative expression (row-wise Z-score of log₂ fold change) of conserved upregulated genes across human, mouse, rat, and zebrafish. Rows represent genes and columns represent species

Hierarchical clustering of these conserved genes across species **(Fig. 4B)** resolved distinct clusters of genes with similar expression profiles thereby highlighting species-specific differences in the magnitude of transcriptional induction. Despite these quantitative differences, the conserved genes showed broadly similar activation patterns across mammals and zebrafish, supporting a shared molecular programme underlying the acute response to brain injury.

Repeating the analysis on the mouse-reference ortholog dataset reproduced these conclusions: although the specific enriched GO terms differed slightly, the analysis again converged on immune activation, inflammatory responses, leukocyte migration, and chemotaxis **(Fig. S3A)**, and hierarchical clustering of the conserved genes yielded comparable cross-species expression patterns **(Fig. S3B)**. The principal biological conclusions remain consistent, identifying immune activation, inflammatory responses, and cell-cycle regulation as conserved early injury responses across the vertebrates irrespective of the ortholog reference employed.

### 3.5 Identification of zebrafish-enhanced regenerative gene signatures following acute TBI

Having established a conserved injury response across vertebrates, we next asked what distinguishes successful regeneration in zebrafish from the regeneration-limited mammals. We therefore filtered for DEGs exhibiting a preferential transcriptional response in zebrafish relative to mammals, defining a gene as zebrafish-enhanced if it was significantly upregulated in zebrafish (log₂FC ≥ 1 and adjusted *P* < 0.05) with a higher expression level than its orthologs in human, mouse, and rat. This approach identified 54 zebrafish-enhanced genes, nominating candidate regulators of regeneration-specific responses.

Functional enrichment of these zebrafish-enhanced genes uncovered biological processes central to tissue repair, immune regulation, and regenerative signalling **(Fig. 5A)**. The most significantly enriched processes included type II interferon-mediated signalling, cellular response to type II interferon, interferon-mediated signalling, cell surface receptor signalling via JAK–STAT and STAT pathways, wound healing, and integrin-mediated signalling, together with myeloid cell differentiation, dendritic cell differentiation, and regulation of erythroid homeostasis, indicating coordinated activation of immune and cellular differentiation programmes. Regeneration-competent zebrafish thus preferentially activate signalling pathways governing inflammatory regulation, extracellular matrix remodelling, and tissue repair after acute brain injury.

**Fig. 5.**
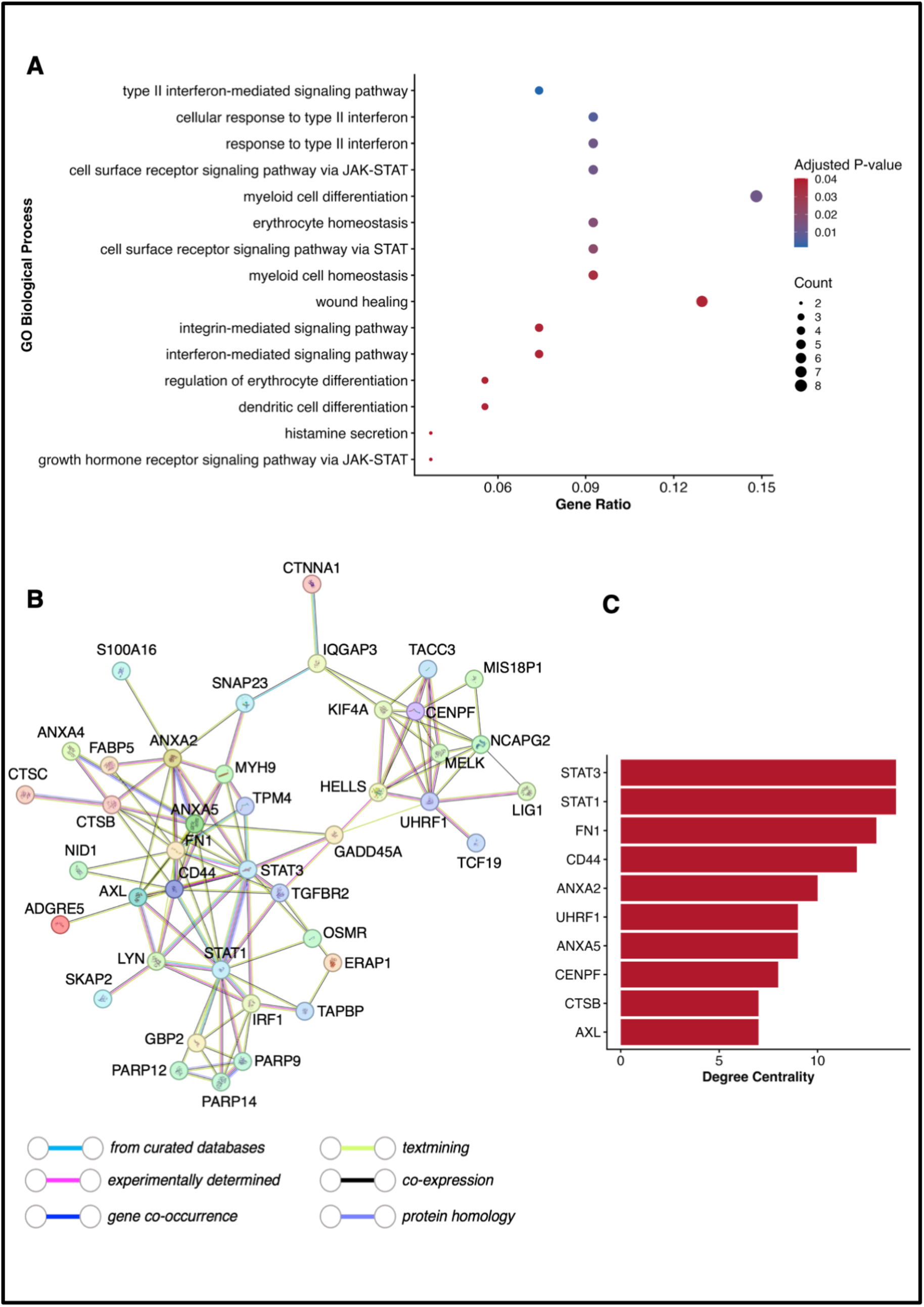
Legend: Identification and network characterization of zebrafish-enhanced regenerative genes following acute traumatic brain injury. (A) Gene Ontology Biological Process enrichment analysis of genes preferentially upregulated in zebrafish compared with mammals, highlighting enrichment of interferon signalling, JAK–STAT signalling, wound healing, integrin-mediated signalling, and cellular differentiation pathways;(B) Protein– protein interaction (PPI) network of zebrafish-enhanced genes constructed using the STRING database, illustrating functional interactions among regeneration-associated proteins;(C) Top ten hub genes identified from the PPI network based on degree centrality, highlighting highly connected regulatory genes including *STAT1*, *STAT3*, *FN1* and *CD44*

We next mapped the functional interactions among these zebrafish-enhanced genes by constructing a protein–protein interaction (PPI) network in the STRING database **(Fig. 5B)**. The network showed extensive connectivity among immune-, signalling-, and extracellular matrix-associated proteins, indicating that these genes act as coordinated regulatory modules rather than isolated molecular events.

Network topology analysis identified several highly connected hub genes by degree centrality: *STAT1* and *STAT3* emerged as the most connected nodes, followed by *FN1*, *CD44*, *ANXA2*, *ANXA5*, *UHRF1*, *CENPF*, *AXL*, and *CTSB* **(Fig. 5C)**. These hub genes converge on cytokine signalling, extracellular matrix remodelling, cell adhesion, and regulation of cellular proliferation, indicating that coordinated activation of these molecular networks may drive the regenerative response observed in zebrafish after TBI.

## 4. Discussion

Traumatic brain injury (TBI) initiates a cascade of molecular events that determines whether damaged neural tissue degenerates or regenerates [42–46]. Adult mammals retain only limited capacity for central nervous system (CNS) repair [47,48], whereas zebrafish restore neural architecture and function with high fidelity [32,49–51], a divergence whose molecular basis remains incompletely understood. By comparing acute-phase transcriptomes across human, mouse, rat, and zebrafish within a unified analytical framework, this study resolves the acute TBI response into a conserved injury programme common to all four species and a regeneration-associated programme unique to zebrafish.

The conserved programme comprised genes governing cell-cycle progression, chromosome segregation, mitotic division, and interferon-mediated immune signalling, indicating that the earliest transcriptional response to CNS injury is evolutionarily conserved irrespective of regenerative outcome. Cell-cycle activation illustrates this duality: in mammals, it drives proliferation of reactive astrocytes, infiltrating immune cells, and vascular endothelial cells that ultimately contribute to glial scar formation [57–60], whereas in zebrafish the same proliferative response precedes expansion of neural progenitor populations that generate replaced neurons [32,49,61]. Interferon and JAK–STAT signalling follow a comparable pattern: both are activated early during the acute neuroinflammatory response [37,62–64] and regulate innate immunity, cytokine signalling, and cellular stress responses [37,62,63,65,66], yet their downstream consequences diverge according to the quality and duration of signalling rather than its presence [64,67–70]. Prolonged inflammatory signalling in mammals is associated with secondary neuronal injury [71,72], whereas zebrafish resolve inflammation rapidly in a manner permissive for neural stem-cell activation [30,56]. These conserved pathways therefore constitute a general injury-response programme rather than a regeneration-specific signature.

Superimposing on this conserved programme, zebrafish selectively activate a second, regeneration-associated transcriptional response. Wound healing was the most prominent zebrafish-enhanced process, consistent with a transition from inflammation to tissue repair: whereas mammalian CNS injury frequently resolves into fibrotic scarring that impedes axonal regeneration [73–77], zebrafish undergo rapid tissue remodelling with minimal fibrosis, sustaining neural progenitor activity [30,56,78,79]. Integrin-mediated signalling reinforces this permissive microenvironment, coordinating cell adhesion and migration during tissue reconstruction [80–83]; dynamic ECM remodelling supports progenitor migration and circuit repair [84–86], and ECM composition itself shapes neural stem-cell behaviour and axonal growth [85–88]. Interferon and JAK–STAT signalling re-emerge within this programme but with regeneration-specific consequences: JAK–STAT signalling governs neural stem-cell activation and proliferation after CNS injury [89,90], and STAT3 activity is specifically required for regenerative neurogenesis in zebrafish [91–93], while interferon signalling additionally regulates progenitor-cell behaviour and tissue repair [94–96]. Coordinated shifts in myeloid and dendritic cell differentiation and erythroid homeostasis [97–99] indicate that zebrafish macrophages and microglia adopt pro-regenerative phenotypes after injury, in contrast to the persistent immune activation associated with secondary damage in mammals [79,100–102].

Protein–protein interaction analysis of the zebrafish-enhanced gene set revealed a highly interconnected regulatory network, with degree centrality identifying *STAT1*, *STAT3*, *FN1*, *CD44*, *ANXA2*, *ANXA5*, *UHRF1*, *CENPF*, *AXL*, and *CTSB* as hub genes. *STAT1* and *STAT3* occupied the most central positions, consistent with their established role as master regulators of cytokine signalling [103,104]; STAT3 activation is specifically required for injury-induced neural stem-cell proliferation and regenerative neurogenesis in zebrafish [56,91,93], whereas dysregulated or sustained STAT signalling in mammals is associated with persistent neuroinflammation and reactive gliosis [29,105,106]. *FN1* and *CD44* formed a second hub centred on ECM regulation: fibronectin mediates cell adhesion, migration, and tissue remodelling [107–109], and CD44 mediates cell–ECM interactions [110,111]; in zebrafish, this remodelling proceeds without the excessive matrix deposition that inhibits axonal regeneration in mammals [112–115]. *AXL*, *ANXA2*, *ANXA5*, and *CTSB* constitute a third functional cluster implicated in immune regulation and membrane repair: AXL regulates phagocytosis and inflammatory signalling after CNS injury [116,117], annexins mediate membrane repair and inflammation resolution [118,119], and cathepsin B contributes to lysosomal processing and ECM turnover [120,121]. *UHRF1* and *CENPF*, associated with DNA replication and chromatin regulation [122–124], remain functionally uncharacterized in the context of zebrafish CNS regeneration and represent priority candidates for future investigation. Collectively, this network topology indicates that regenerative competence arises from coordinated activity across this gene set rather than from any single regulatory node.

These findings indicate that regenerative competence following TBI is not determined by activation of a single regeneration-associated gene, but by the layering of an additional, zebrafish-specific transcriptional programme onto a conserved injury response, consistent with evidence that zebrafish CNS regeneration depends on coordinated interactions between neural stem cells, immune cells, and extracellular signalling [30,32,52–56]. Mammals and zebrafish therefore initiate a similar early transcriptional response to CNS injury; regenerative failure in mammals may instead reflect the absence of mechanisms required to convert this conserved response into productive tissue repair.

Several limitations should be noted. The included datasets derive from heterogeneous laboratories, sequencing platforms, and injury models; although REML-based meta-analysis and standardized pre-processing reduced technical variability, residual heterogeneity cannot be excluded. This analysis was restricted to the acute phase of TBI, and the molecular programmes governing subsequent stages of neurogenesis and functional recovery remain uncharacterized. Because the data are transcriptomic, the hub genes identified here represent statistical rather than causal regulators of regeneration; functional validation using gene knockdown, CRISPR-mediated perturbation, or transgenic zebrafish models will be required to establish causality. Ortholog mapping, finally, does not fully account for zebrafish-specific gene duplication arising from teleost-specific whole-genome duplication, which may obscure additional regeneration-associated mechanisms unique to this lineage.

## 5. Conclusion

This study provides a cross-species transcriptomic framework that resolves the acute response to traumatic brain injury into a conserved injury programme shared by human, mouse, rat, and zebrafish and defined by cell-cycle activation and interferon-mediated signalling and a pro-regenerative programme unique to zebrafish, comprising wound healing, ECM remodelling, and JAK–STAT signalling, coordinated through a hub-gene network centred on *STAT1*, *STAT3*, *FN1*, *CD44*, and *AXL*.

These findings indicate that regenerative competence reflects the coordinated regulation of multiple interacting molecular programmes rather than the action of a single gene, and that the mammalian CNS retains much of the conserved injury response required for regeneration. Defining the mechanisms that enable zebrafish to convert this conserved response into productive repair may inform strategies to enhance endogenous CNS repair after traumatic brain injury in humans.

## Abbreviations

TBI: Traumatic brain injury
CNS: central nervous system
REML: restricted maximum likelihood
RNA-seq: bulk RNA sequencing
scRNA-seq: single-cell RNA sequencing
GEO: Gene Expression Omnibus
DEGs: differentially expressed genes
GO: Gene Ontology;

## Author Contributions

PG and SPH conceptualised and designed the study. PG collected data and performed the analysis. PG and SPH wrote the manuscript. SPH supervised all aspects of the study.

## Data availability statement

This study utilises publicly accessible data extracted from published studies; therefore, no original data are available for sharing beyond the extracted dataset summarised in Supplementary Tables, which is available from the corresponding author upon reasonable request.

## Declaration of generative AI use

During preparation of this manuscript, the authors used AI-assisted language-editing tools solely to improve clarity of expression. All scientific content, data, analyses, and interpretations are the authors’ own. The authors reviewed and edited the output as needed and take full responsibility for the content of the publication.

