## Supplementary Figures for "Comparative transcriptomic analysis reveals conserved injury responses and divergent molecular responses underlying early regenerative competence following traumatic brain injury"

**Supplementary Figure S1**

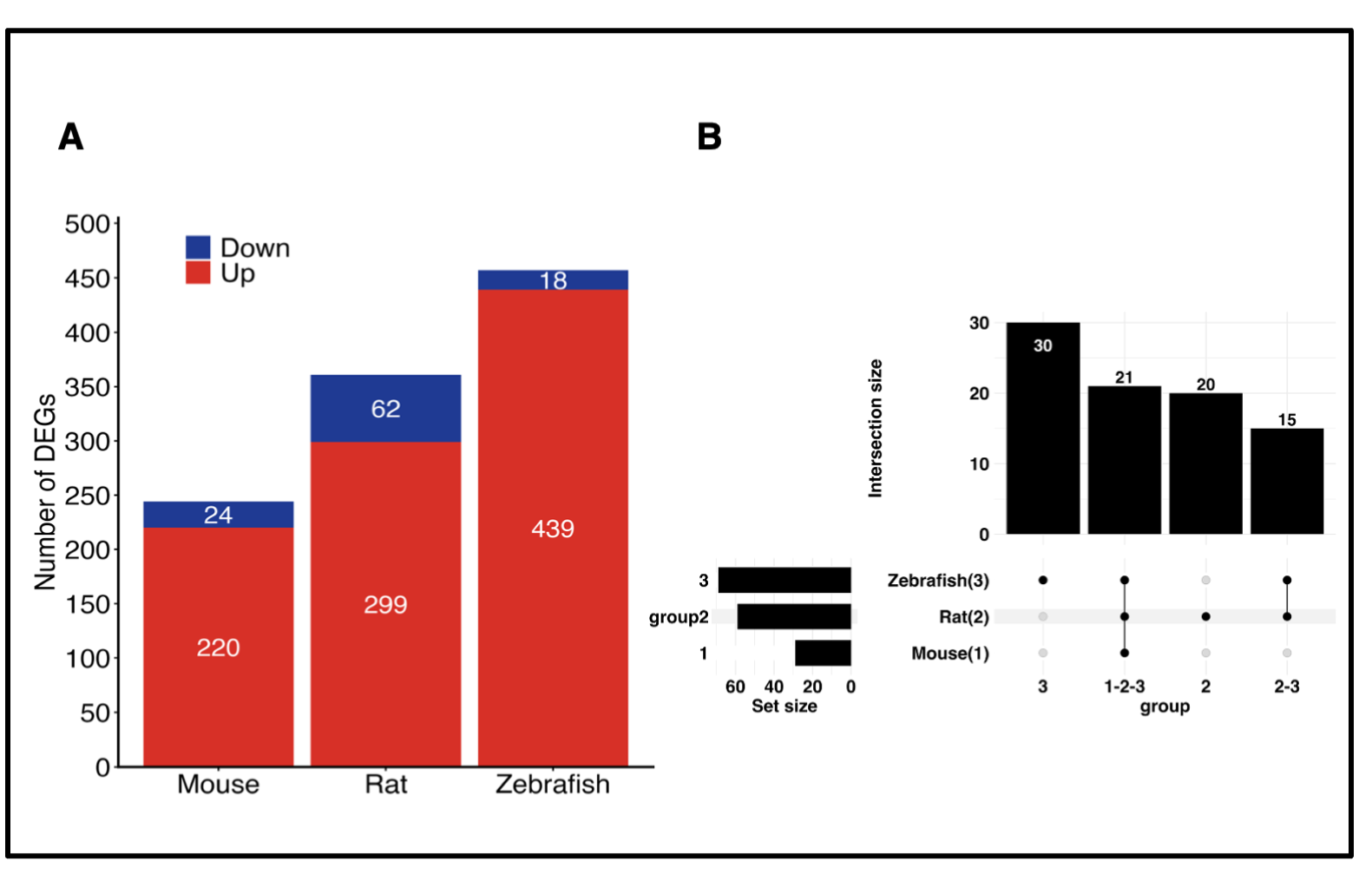

**Supplementary Figure S1 Legend: Global differential gene expression patterns in the mouse-reference comparative transcriptomic dataset following acute traumatic brain injury.**

(A) Number of significantly differentially expressed genes (DEGs) identified in the mouse-reference dataset comprising mouse, rat, and zebrafish following species-specific differential expression analysis and REML-based integration. Differentially expressed genes were defined using an absolute log₂ fold change (|log₂FC|) ≥ 1 and an adjusted *P* value < 0.05. Red bars represent upregulated genes, whereas blue bars represent downregulated genes. Zebrafish exhibited the largest number of DEGs, followed by rat and mouse, with transcriptional activation predominating in all three species.

(B) UpSet plot showing the overlap of orthologous DEGs among mouse, rat, and zebrafish using the mouse-reference comparative dataset. Horizontal bars represent the total number of DEGs identified in each species (set size), whereas the vertical bars indicate the number of genes shared among different combinations of species (intersection size). Filled black circles connected by vertical lines denote the species included in each intersection. Similar to the human-reference analysis, most DEGs were species-specific or shared among a limited number of species, whereas a smaller subset of genes was conserved across all three species, demonstrating the robustness of the comparative transcriptomic framework.

**Supplementary Figure S2**

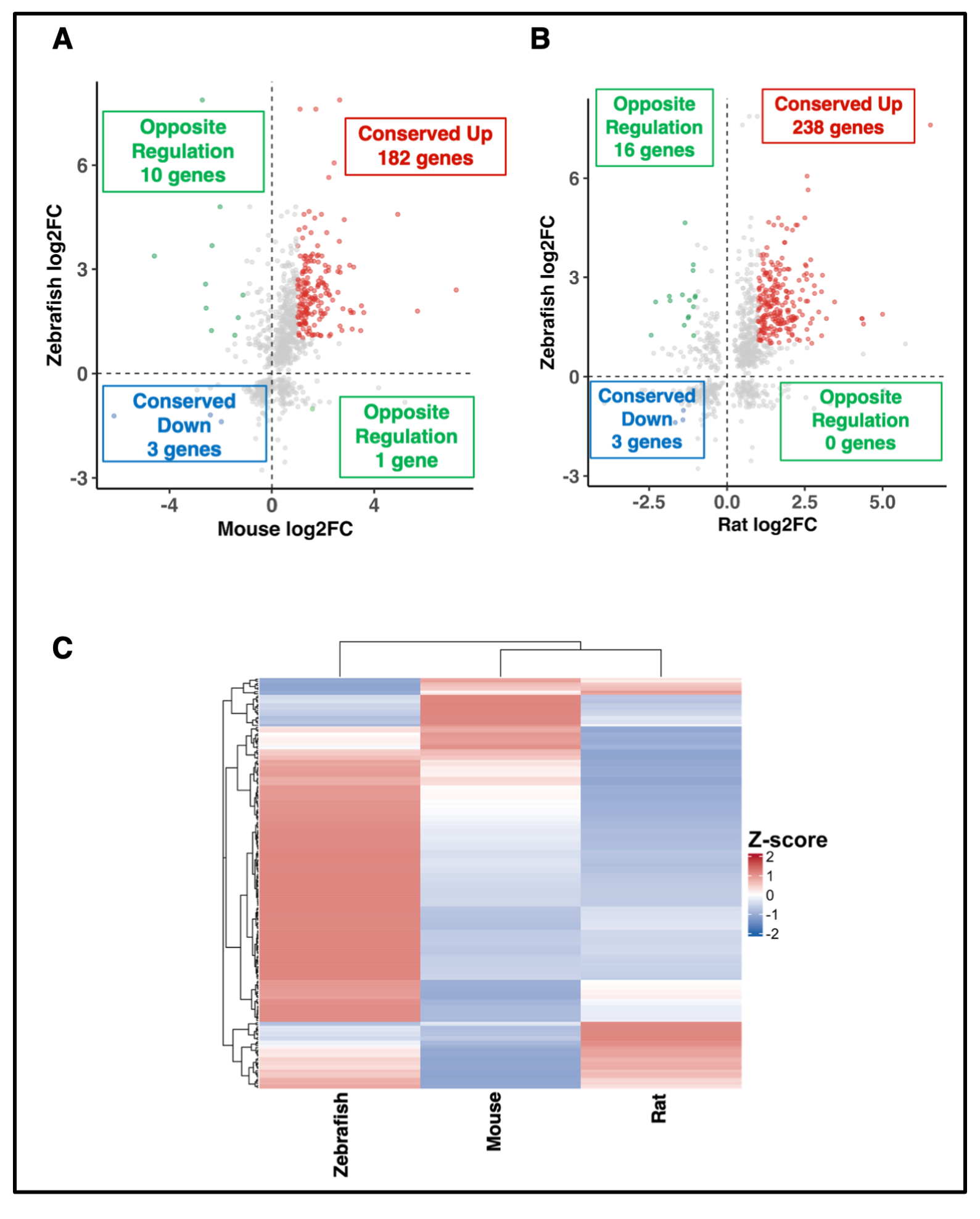

**Supplementary Figure S2 Legend: Comparative transcriptomic analysis among regeneration-competent species using the mouse ortholog reference dataset.**
(A) Scatter plot comparing differential gene expression between mouse and zebrafish following acute traumatic brain injury (TBI). Each point represents an orthologous gene, plotted according to its meta-analysis log₂ fold change (log₂FC) in mouse (x-axis) and zebrafish (y-axis). Genes exhibiting conserved upregulation (upper-right quadrant; *n* = 182) and conserved downregulation (lower-left quadrant; *n* = 3) are highlighted in red and blue, respectively, whereas oppositely regulated genes are shown in green (upper-left, *n* = 10; lower-right, *n* = 1). Gray points represent genes not meeting the differential expression criteria in both species.

(B) Scatter plot comparing rat and zebrafish differential gene expression using mouse-reference ortholog mapping. Conserved upregulated genes (upper-right quadrant; *n* = 238), conserved downregulated genes (lower-left quadrant; *n* = 3), and oppositely regulated genes (upper-left, *n* = 16; lower-right, *n* = 0) are indicated. Gray points denote genes outside the conserved or opposite regulation categories.

(C) Hierarchical clustering heatmap of conserved differentially expressed genes identified across zebrafish, mouse, and rat. Rows represent orthologous genes, columns represent species, and colours indicate row-wise Z-score normalized log₂FC values (red, relatively higher expression; blue, relatively lower expression). Dendrograms illustrate similarity in species-specific expression profiles and gene clustering.

**Supplementary Figure S3**

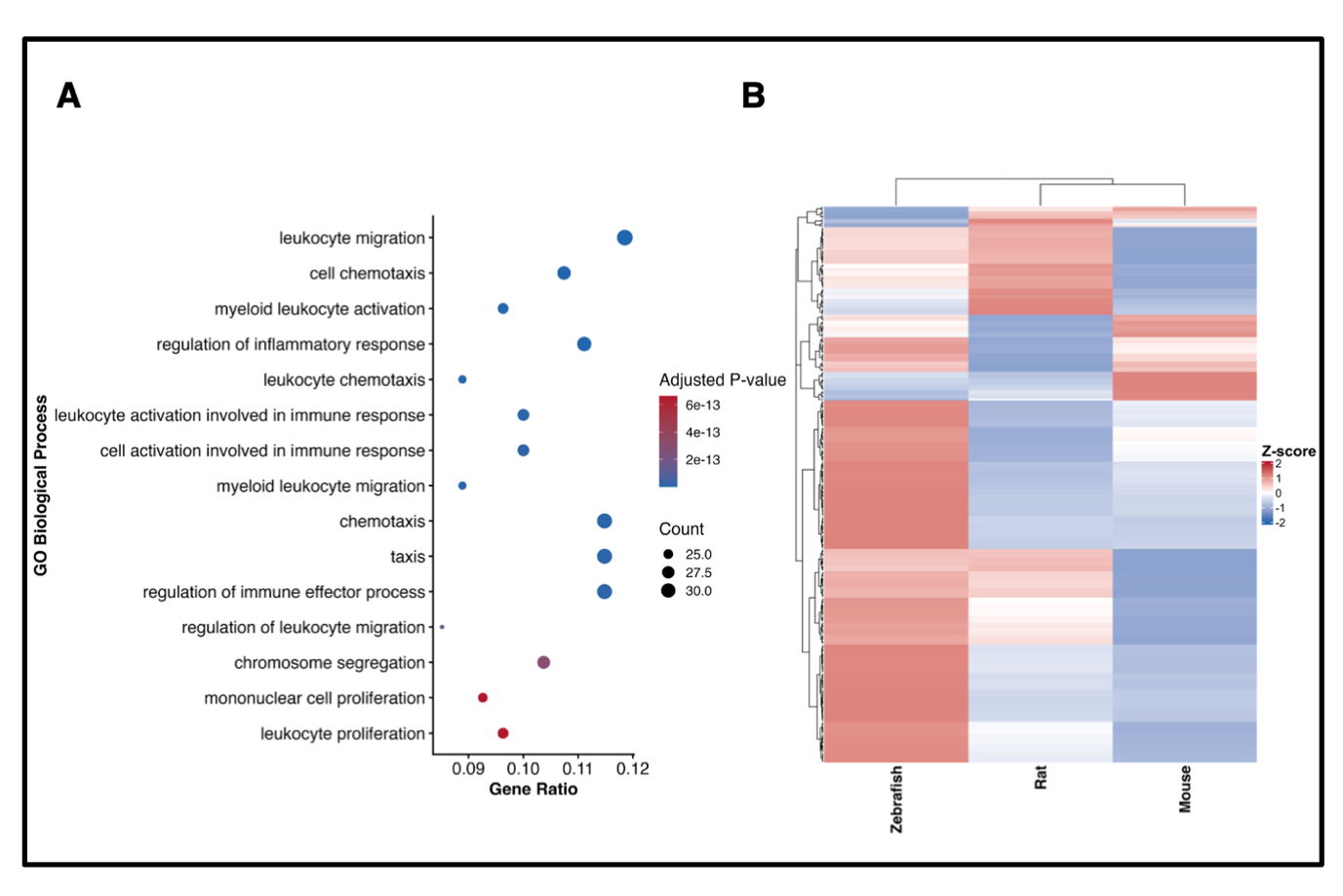

**Supplementary Figure S3 Legend: Functional enrichment analysis using the mouse ortholog reference dataset.**

(A) GO Biological Process enrichment analysis of conserved upregulated genes identified using mouse-reference ortholog mapping.

(B) Hierarchical clustering heatmap showing the relative expression (row-wise Z-score of log₂ fold change) of conserved genes across mouse, rat, and zebrafish, with mapped human orthologs where applicable. The analysis demonstrates that the major biological conclusions are robust to the choice of ortholog reference.
